# Adolescent brain-age differences between profiles based on suicidal ideation, distress and wellbeing

**DOI:** 10.64898/2026.07.31.741956

**Authors:** Maddison Crethar, Laura K M Han, Taliah Prince, Lia Mills, Timothy J Silk, Nandita Vijayakumar, Daniel F. Hermens, Amanda Boyes

**Affiliations:** Thompson Institute, University of the Sunshine Coast, Birtinya, Queensland, Australia; Department of Psychiatry, Amsterdam UMC location Vrije Universiteit Amsterdam, Amsterdam, The Netherlands; Amsterdam Neuroscience, Mood, Anxiety, Psychosis, Sleep & Stress program, Amsterdam, The Netherlands; Amsterdam Public Health, Mental Health program, Amsterdam, The Netherlands; Deakin Lifespan Institute and School of Psychology, Deakin University, Melbourne, Australia; Developmental Imaging, Murdoch Children’s Research Institute, Parkville, Victoria, Australia; Centre for Adolescent Health, Murdoch Children’s Research Institute, Parkville, Victoria, Australia

**Keywords:** Brain-age, adolescence, suicidality, structural MRI, psychological distress, wellbeing

## Abstract

**Background:** Predicted brain-age derived from structural neuroimaging is being increasingly explored as a marker of brain maturation and biological age, with mental ill-health linked to advanced biological ageing in adults. Longitudinal evidence in adolescents remains limited, and the relationship between suicidality, psychological distress, wellbeing and brain-age gap (BrainAGE; predicted brain-age minus chronological age) is poorly understood.

**Methods:** Data were drawn from 135 adolescents (54.8% female; 12-16.9 years; n=688 observations) from the Longitudinal Adolescent Brain Study. Latent profile analysis (LPA) used person-level means and standard deviations of distress (K10), wellbeing (COMPAS-W) and suicidal ideation (SIDAS). BrainAGE was estimated using the CentileBrain Global-BrainAGE pipeline from FreeSurfer-derived morphometric features. Associations between cluster membership and BrainAGE were examined using linear regression and linear mixed-effects models, adjusted for chronological age and sex.

**Results:** Three profiles emerged: low distress (N=110; 50% female), moderate distress with greater suicidal ideation (N=14; 64% female) and moderate distress with lower suicidal ideation (N=11; 91% female). The moderate distress with lower suicidal ideation cluster showed significantly higher BrainAGE relative to the low distress cluster (B=1.87, SE=0.76, p=.015). Whereas, the moderate distress with greater suicidal ideation cluster did not differ to the low distress cluster. Chronological age was positively associated with BrainAGE (B=0.67, SE=0.07, p<.001).

**Conclusions:** Distinct adolescent mental health profiles may be associated with BrainAGE, depending on the levels and stability of distress and suicidality. However, the longitudinal associations were less robust across small extensions of the developmental range studied, warranting some cautious interpretation and need for replication.

## 1. Introduction

Chronological age reflects time since birth; however, it only partially reflects health and functional characteristics (1). In contrast, predicted age metrics are used to study health-related trajectories (1) and may be biological, functional or phenotypic (2). Within this framework, imaging-derived brain-based predicted brain-age (PBA) provides an estimate of biological age from structural neuroimaging data and has been used to examine deviations from typical brain development and ageing (3–6). Although ageing biomarkers exhibit substantial inter-individual variability due to a range of biopsychosocial factors, they are generally considered robust for group-level comparisons (4). The deviation between PBA and chronological age is inherently a linear construct, representing either a positive or negative brain-age gap (BrainAGE). However, brain structural changes may exhibit nonlinear trajectories across childhood, adolescence, and young adulthood, reflecting the rapid and heterogeneous patterns of maturation (7). Environmental factors such as socioeconomic disadvantage may further shape these trajectories, with mixed evidence on their association with PBA (8, 9).

A growing body of evidence links stress and mental ill-health to advanced biological ageing, although most evidence has emerged from cross-sectional studies and adult cohorts (7, 10). Mental disorders, particularly when comorbid, have been associated with more “advanced” biological ageing (10), and are thought to promote age-related alterations via chronic inflammation (11), dysregulation of the hypothalamic-pituitary-adrenal (HPA) axis (12), and mitochondrial dysfunction and epigenetic modifications (3). These associations may be especially relevant in adolescence, a period of rapid cortical, hormonal and psychosocial change (13–16), in which stress exposure has been linked to accelerated pubertal development and advanced brain-ageing, although findings vary by sex, development stage and context (9, 17, 18). Environmental stressors may shape neurodevelopmental trajectories, with neighbourhood disadvantage in early adolescence associated with advanced brain-ageing (8). Psychological stress can also alter brain structure and function with adverse childhood experiences associated with changes in brain networks and microstructure, with differences according to biological sex and pubertal timing (19–23).

In adolescents, psychological distress is a well-established risk factor for suicidality, but its relationship with BrainAGE remains poorly understood. Research examining BrainAGE in young people has primarily focused on broad psychiatric diagnoses and transdiagnostic mental health risk (24, 25). Importantly, other psychological constructs and behaviours may relate independently to BrainAGE. Specifically, while often associated with distress and depressive symptoms, mental wellbeing and suicidal ideation (SI) can be conceptually and empirically distinct (26–29). Hence, identifying brain-ageing patterns across adolescent profiles of distress, suicidality and wellbeing may improve understanding of mental health risk and inform early intervention, particularly given evidence that repeated measures improve the reliability of biological age indicators (7) and that advanced ageing is associated with later health burden (10, 30). The present study, therefore, examined whether person-centred mental health profiles differ in mean BrainAGE and predicted brain-ageing trajectories, across adolescence, hypothesising higher BrainAGE among adolescents with greater suicidality and psychological distress, and lower wellbeing.

## 2.0 Methods and Materials

### 2.1 Study Design and Participants

This study utilised data from the ongoing Longitudinal Adolescent Brain Study (LABS), collected between 18 July 2018 and 25 February 2025. LABS design and recruitment details have been described previously and can be found elsewhere (31–33). Ethics approval (A181064) was granted by the University of the Sunshine Coast Research Ethics Committee. Written consent was obtained from participants and parents/caregivers. Assessments took up to five hours to complete, with breaks scheduled, and all were completed with trained researchers at UniSC’s Thompson Institute.

Selection criteria for LABS required participants to be aged 12-15 years at enrolment and proficient in spoken and written English. An additional inclusion criterion specific to this study was that participants must have attended at least two timepoints to contribute to longitudinal analyses. Young people were excluded from the study if they had suffered a major neurological disorder or major medical illness, had a sustained head injury (with loss of consciousness >30 minutes), had an intellectual disability, or if they were unable to complete an MRI. The final sample included 135 participants with 688 observations. The sample reduced following initial MR quality checks (n=147 scans were excluded due to scan quality [i.e., artefact or bad motion]). Consistent with recommendations by Boyes et al. (31), analyses were restricted to observations collected between ages 12 and 16 years to reduce developmental variability, resulting in a sample of 136 participants. One additional participant was excluded for having fewer than two timepoints with MRI and self-report data, resulting in a final dataset of N=135.

### 2.2 Self-Report Measures

As part of a larger self-report questionnaire, participants completed the following scales:

#### Psychological Distress

The 10-item Kessler psychological distress (K10) scale (34) with items answered on a 5-point scale (1 = *none of the time* to 5 = *all of the time*). Overall scores ranged from 10-50 (higher scores = greater psychological distress). The K10 is a valid measure of recent psychological distress in adolescents, which uses the participant’s total score to identify their level of non-specific psychological distress and likelihood of psychological disorders within the last 30 days (34, 35).

#### Wellbeing

The 26-item COMPAS-W scale measures psychological and subjective wellbeing (36) with items answered on a 5-point Likert-type scale (1 = *strongly disagree* to 5 = *strongly agree*). There are three measures of eudaimonic (psychological) wellbeing (‘own-worth’, ‘mastery’ and ‘achievement’) and three measures of hedonic (subjective) wellbeing (‘composure’, ‘positive’ and ‘satisfaction’). Overall wellbeing scores range from 26-130, with higher scores indicating greater self-reported wellbeing (36, 37).

#### Suicidality

The five-item Suicidal Ideation Attributes Scale (SIDAS) is designed to identify suicidal thoughts and their severity (38). Responses are rated on an 11-point Likert scale (0 = *never,* to 10 = *always*). Total scores range from 0 to 50 (higher scores = greater severity of suicidal ideation) (39). The SIDAS has high validity and reliability (38) and has been validated for use with 16–25-year-olds (40). Participants also completed four items relating to suicidal thoughts and behaviours from the Youth Risk Behaviour Survey (YRBS) which assess the presence of suicidal thoughts and behaviours within the past 12 months. Scores for items 1 and 2 range from 0 to 1 (0 = *no*, 1 = *yes*). Scores on item 3 range from 0 to 4 (0 = *0 times*, to 4 = *6 or more times*). Lastly, scores on item 4 range from 0 to 1 (0 = I did not attempt or no, 1 = yes). Total scores are calculated as a sum of each question, with higher scores indicating greater severity. The YRBS suicidality items have demonstrated good convergent and discriminant validity in adolescents (41).

### 2.3 Magnetic Resonance Imaging acquisition and processing

Structural MRI brain scans were acquired using a 3-Tesla Siemens Skyra scanner (Erlangen, Germany) with a 64-channel head and neck coil, at the Nola Thompson Centre for Advanced Imaging (Thompson Institute, UniSC) in Birtinya, Australia. The MRI protocol included a T1-weighted magnetisation prepared rapid acquisition gradient echo sequence (MPRAGE; TR=2200ms, TE=1.71ms, TI=850ms, flip angle=7°, spatial resolution=0.9x0.89x0.89mm, FOV=208x256x256, TA=3:57). Data was extracted via the FreeSurfer (7.4.0) longitudinal recon-all pipeline (Reuter et al., 2012), packaged in singularityCE (version 3.8.0), on the UniSC HPC system, using PBS (version 20.0.0), Ubuntu 20.04, and connected to a CEPH cluster (version 15.2.13).

#### MR Quality Check

All scans were visually inspected for quality issues (movement or artefact). Following FreeSurfer processing scan quality was further assessed using the ENIGMA FreeSurfer Cortical & Subcortical Extraction and QC Protocol (available: https://github.com/ENIGMA-git/ENIGMA-FreeSurfer-protocol; Hibar et al., 2015), and Euler numbers. Outliers determined via ENIGMA QC protocol and scans with Euler Z scores of ±3.0 underwent further (cortical and subcortical) segmentation inspection.

#### Brain-age Calculations and Processing

Adjusted Brain-age Gap (BrainAGE) was calculated using the CentileBrain Global-BrainAGE model. This model is an open-science, web-based platform (https://centilebrain.org/#/brainAge_global) (6, 42) that uses 150 morphometric features, including 68 cortical thickness measures, 68 cortical surface area measures and 14 subcortical volumes. These features were extracted from FreeSurfer outputs, with cortical measures obtained from the Desikan-Killiany atlas parcellation (aparc) and subcortical volumes extracted from the aseg segmentation. The 5≤age≤40 years model was utilised to extract adjusted BrainAGE (predicted ‘brain-age’ minus chronological age adjusted to remove dependence on chronological age) output by sex (6, 42).

### 2.4 Statistical Analyses

The following analyses were conducted in R Studio version 4.4.1.

#### Latent Profile Analysis

was conducted using the *tidyLPA* package (version 2.0.2), implementing Gaussian mixture modelling with equal variances and covariances fixed to zero across clusters (13). The dataset was restructured from long format to person-level format (one row per participant). This was done by computing the mean score and standard deviation (SD) across all available timepoints for each participant on three self-report scales (COMPAS-W, K10 and SIDAS). This created six LPA indicators per participant, with three mean scores capturing each individual’s overall level of mental health across the study period and three SD values capturing the degree to which scores fluctuated across timepoints attended by that participant. All six indicators were reviewed for outliers. Values were winsorised at the 2.5^th^ and 97.5^th^ percentiles (i.e. values outside of these boundaries were capped at the respective percentile). The number of outliers did not exceed 10% for any variable. Following winsorisation, all indicators were standardised to z-scores to ensure equal weighting across scales.

Models with one, two and three cluster solutions for the LPA were estimated and compared on the log-likelihood (43), Akaike information criterion (AIC) (44), Bayesian information criterion (BIC) (45), sample size-adjusted BIC (SABIC) (46), entropy, and bootstrapped likelihood ratio test (BLRT) (47). Model selection prioritised lower BIC and SABIC values, high entropy (indicating acceptable class separation), a significant BLRT (indicating meaningful improvement over the previous solution) (43), and a minimum profile size of 5% of the analytic sample (48).

#### Brain-age analysis

Two complementary models were examined to explore whether cluster membership was associated with BrainAGE, using the *lme4* and *lmerTest* packages. First, a person-level linear regression was conducted with mean BrainAGE (averaged across all available timepoints per participant) as the outcome, cluster membership as the predictor (Cluster 1 was set as the reference), and chronological age and biological sex (female, male) as covariates (mean_BrainAGE ∼ cluster + sex + mean_age). Then, a secondary longitudinal analysis was completed on the entire dataset (688 observations) using a linear mixed-effects model (BrainAGE ∼ cluster + sex + age + (1 | Participant). BrainAGE at each timepoint was the outcome, cluster membership, chronological age and sex were fixed effects, with a random intercept for participant to account for the non-independence of repeated observations (at 4-monthly intervals).

#### Post hoc analysis

Three additional models were run as diagnostic tests utilising the entire dataset: (i) the secondary longitudinal model this time without the random intercept for participant, treating repeated observations as independent to demonstrate the effect of ignoring within-person grouping on standard errors and significance (BrainAGE ∼ cluster + sex + age); (ii) the secondary longitudinal model without sex as a covariate (as the Centile BrainAGE pipeline produces sex specific estimate); BrainAGE ∼ cluster + age + (1 | ParticipantID)); (iii) the secondary longitudinal model for the 12-17.9 dataset to examine the influence of including older participants with greater variation in BrainAGE, as detected via the methods analysis described in Boyes et al. (31).

## 3. Results

### 3.1 LPA results

LPA model fit comparisons across the one-to three-cluster solutions are presented in Table 1. A three-cluster solution was selected based on having the lowest AIC, BIC and SABIC values, significant BLRT, high entropy, and interpretability (43–45, 47). Cluster 1 had the highest participant count (*N*=110; 50.0% female) and is labelled ‘low distress’ based on mean scores (and lowest deviations). Specifically, participants reported the highest levels of wellbeing (COMPAS-W; M=99.54; SD=5.56), the lowest psychological distress (K10; M=16.20; SD=2.81) and lowest SI (SIDAS; M=0.19; SD=0.39). Cluster 2 (*N*=14; 64.3% female) was labelled as ‘moderate distress with greater SI’, as this subgroup reported the lowest levels of wellbeing (COMPAS-W; M=85.93; SD=6.89) and highest levels of psychological distress (K10; M=26.41; SD=6.07) and SI (SIDAS: M=10.82; SD=10.96). Cluster 3 (N = 11; 90.9% female), labelled as ‘moderate distress with lower SI’ had moderate levels of wellbeing (COMPAS-W; M=93.12; SD=6.82), psychological distress (K10; M=22.89; SD=5.80) and SI (SIDAS: M=4.31; SD=6.75). Table 2 summarises demographic details as well as suicidal thoughts and behaviours across clusters. Standardised mean scores and standard deviation stability of the three cluster groups are depicted in Figure 1, with cluster group means and individual trajectories of participants shown in Figure 2.

**Figure 1.**
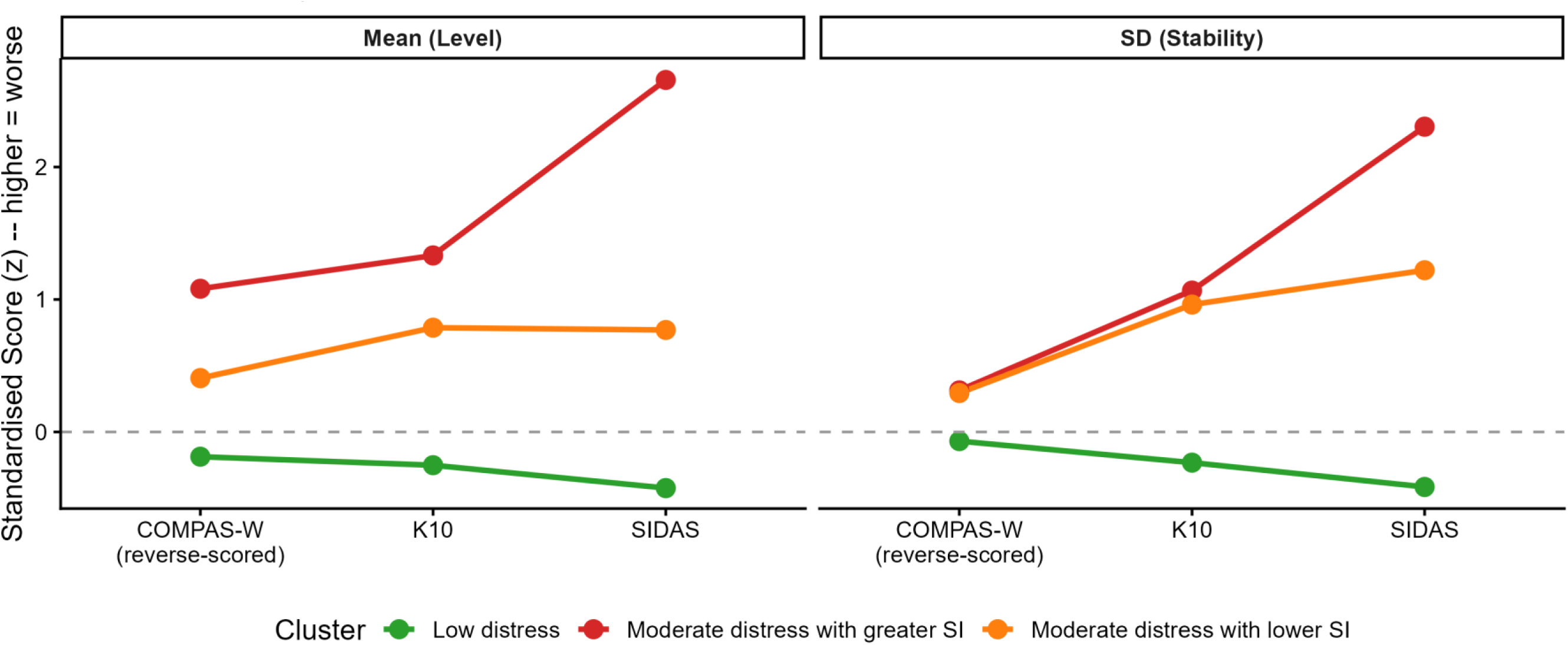
The average and stability of measures included in the LPA determined profiles (3-Cluster solution) COMPAS-W = wellbeing; K10 = psychological distress; SIDAS = suicidal ideation; COMPAS-W was reverse-scored so that a higher score represents a worse score, in line with K10 and SIDAS.

**Figure 2.**
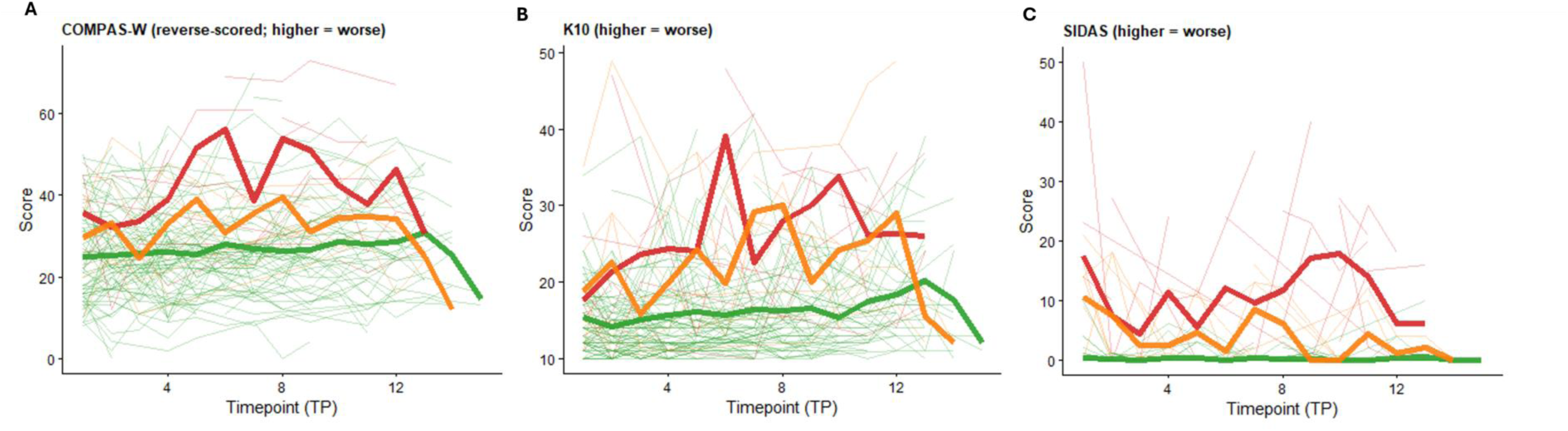
Individual and mean trajectories according to cluster groups. (A) COMPAS-W = wellbeing; (B) K10 = psychological distress; (C) SIDAS = suicidal ideation; Green = low distress; Red = moderate distress with greater SI; Orange = moderate distress with lower SI.

**Table 1.** Model fit comparisons across 1-3 cluster solutions. AIC = Akaike information criterion; BIC = Bayesian information criterion; SABIC = sample size-adjusted BIC; BLRT = bootstrapped likelihood ratio test.

| Number of clusters | Log-likelihood | AIC | BIC | SABIC | Entropy | BLRT $p$ |
| --- | --- | --- | --- | --- | --- | --- |
| 1 cluster solution | -1146.81 | 2317.63 | 2352.49 | 2314.53 | 1 | NA |
| 2 cluster solution | -945.94 | 1929.88 | 1985.08 | 1924.98 | 0.985 | 0.010 |
| 3 cluster solution | -875.15 | 1802.29 | 1877.83 | 1795.58 | 0.988 | 0.010 |

**Table 2.** Demographic and prevalence of suicidal thoughts and behaviours across clusters for N=135, n=688 datasets. Values represent participant count (sex and socioeconomic status) and participant-level means and standard deviations averaged across timepoints. YRBS = Youth Risk Behavior Survey; SI = suicidal ideation. YRBS items were not included in latent profile estimation and are presented for descriptive purposes only. Higher values indicate greater endorsement of suicidal thoughts and behaviours.

| Item | Low Distress<br>(N=110; n=575) M<br>(SD) | Moderate Distress with<br>Greater SI (N=14; n =<br>51) M (SD) | Moderate Distress with<br>Lower SI (N=11; n =<br>62) M (SD) |
| --- | --- | --- | --- |
| Age | 14.08 (0.99) | 14.69 (1.14) | 14.34 (0.82) |
| **Sex** |  |  |  |
| Female | 55 | 9 | 10 |
| Male | 55 | 5 | 1 |
| **Socioeconomic Status** |  |  |  |
| Quintile 1 <sup>a</sup> | 4 | 1 | 1 |
| Quintile 2 | 3 | 1 | - |
| Quintile 3 | 42 | 3 | 5 |
| Quintile 4 | 55 | 8 | 5 |
| Quintile 5 <sup>b</sup> | 6 | 1 | - |
| **YRBS Items** |  |  |  |
| Considered suicide | 0.01 (0.07) | 0.47 (0.30) | 0.10 (0.15) |
| Made a suicide plan | 0.01 (0.05) | 0.36 (0.34) | 0.04 (0.09) |
| Attempted suicide | 0.00 (0.00) | 0.56 (0.95) | 0.04 (0.12) |
| Injurious suicide attempt | 0.00 (0.00) | 0.04 (0.13) | 0.02 (0.06) |

<sup>a</sup> Most advantaged socioeconomic status quintile
<sup>b</sup> Most disadvantaged socioeconomic status quintile

### 3.2 Linear Regression Analysis

The overall model was statistically significant [*F*(4,130)=5.63, *p*<.001, *R²*=.148 (adjusted *R²=*.122)], indicating that cluster membership, biological sex and chronological age together explained approximately 14.8% of the variance in BrainAGE. As shown in Table 3, the *moderate distress with greater SI* cluster did not differ significantly from *low distress* cluster in BrainAGE (*B*=-0.51, *SE*=0.60, *t*(130)=-0.86, *p*=.396, 95% CI [-1.70, 0.68]). Whereas, the *moderate distress with lower SI* cluster had a significantly higher BrainAGE compared to the *low distress* cluster (*B*=2.01, *SE*=0.67, *t*(130)=2.97, *p*=.004, 95% CI [0.67, 3.34]). Participants in the *moderate distress with lower SI* cluster had a BrainAGE ∼2.01yrs older than those in the *low distress* cluster after controlling for sex and age. Chronological age was also a significant predictor of BrainAGE (*B*=-0.49, *SE*=0.18, *t*(130)=-2.67, *p*=.009, 95% CI [-0.86, - 0.13]), indicating that older participants tended to have higher BrainAGE values. Biological sex was not a statistically significant predictor of BrainAGE (*B*=0.70, *SE*=0.37, *t*(130)=1.86, *p*=.065, 95% CI [-0.04, 1.44]).

**Table 3.** Linear regression of adjusted brain-age gap on cluster membership. The *low distress* cluster served as a reference group. *B* = unstandardised regression coefficient; *SE* = standard error; *t* = t-statistic; *CI* = confidence interval. The model controlled for biological sex and chronological age.

| Predictor | *B* | *SE* | *t* | *p* | 95% CI |
| --- | --- | --- | --- | --- | --- |
| Intercept | 3.86 | 2.58 | 1.50 | 0.136 | [-1.24, 8.97] |
| *Moderate distress with greater SI vs Low distress* | -0.51 | 0.60 | -0.85 | 0.396 | [-1.70, 0.68] |
| *Moderate distress with lower SI vs Low distress* | **2.01** | **0.67** | **2.97** | **0.004** | **[0.67, 3.34]** |
| Sex (female vs male) | 0.70 | 0.37 | 1.86 | 0.065 | [-0.04, 1.44] |
| Chronological age | **-0.49** | **0.18** | **-2.67** | **0.009** | **[-0.86, 0.13]** |

To evaluate the robustness of the primary findings, a linear mixed-effects model was conducted across the entire dataset, with BrainAGE as the outcome, cluster membership, chronological age and biological sex as fixed effects and a random intercept for participant. Consistent with the primary analysis, the *moderate distress with lower SI* cluster had significantly higher BrainAGE than the *low distress* cluster (*B*=1.87, *SE*=0.76, *t*(119.27)=2.48, *p*=.015, 95% CI [0.38, 3.37]). Whereas, the *moderate distress with greater SI* cluster did not differ significantly from the low distress cluster (*B*=-1.19, *SE*=0.68, *t*(133.82)=-1.74, *p*=.083, 95% CI [-2.54, 0.16]). Chronological age was a significant positive predictor of BrainAGE (*B*=0.67, *SE*=0.07, *t*(642.88)=10.05, *p*<.001, 95% CI [0.54, 0.80]), whereas biological sex was not associated with BrainAGE (*B*=0.37, *SE*=0.42, *t*(124.85)=0.88, *p*=.380, 95% CI [-0.46, 1.20]. Figure 3 plots BrainAGE against chronological age for each cluster. All clusters demonstrated positive associations between chronological age and BrainAGE, although the strength of this relationship differed. The *low distress* cluster showed a moderate positive association between chronological age and BrainAGE (*R²*=0.012), the *moderate distress with greater SI* cluster had the weakest relationship (*R²<*.001) and the *moderate distress with lower SI* cluster had the strongest positive relationship between chronological age and BrainAGE (*R²*=0.122).

**Figure 3.**
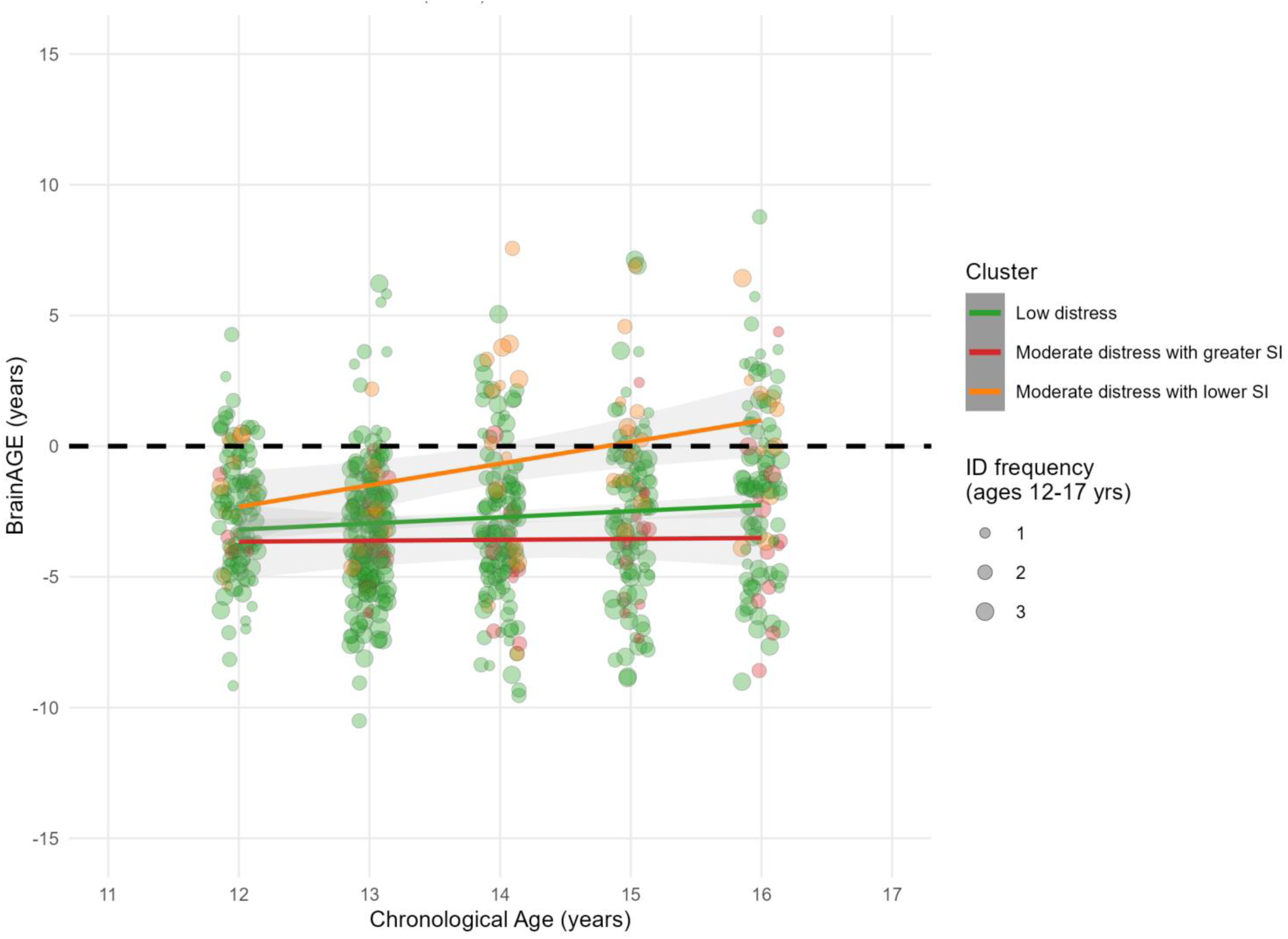
BrainAGE vs chronological age. Points represent individual observations across all assessment timepoints. Point size reflects the frequency of observations within each age-cluster combination. Solid lines represent cluster-specific linear regression fits with 95% confidence intervals. The dashed line indicates the 1:1 reference line at which predicted brain-age equals chronological age (zero brain-age gap).

### 3.3 Post hoc analyses

To demonstrate the sensitivity of statistical significance to the repeated-measures cluster structure, the linear mixed effects model was re-run (i) without participant ID as a random intercept, and (ii) without sex as a covariate. The results are presented in *Supplementary Material S1.* Results revealed that when the random intercept for participant was removed, the direction and magnitude of cluster effects were similar, however two effects that were non-significant in the linear mixed model reached significance in the fixed-effects model (*moderate distress with greater SI*, *p*=.016; sex, *p*=.002). Removing sex from the secondary, longitudinal analysis did not meaningfully change cluster effects or their significance (see *S1)*, supporting the robustness of the cluster findings to this covariate. Lastly, the secondary, longitudinal analysis was re-run on the entire 12-17.9-year dataset (N=137 with a minimum of 2 TPs [MRI, self-report] data; n=750 observations) (see S2, S3). Neither *moderate distress* cluster differed significantly from the *low distress* cluster. Age remained a significant predictor of BrainAGE (*B***=**0.72, *p*<.001).

## 4. Discussion

This study examined BrainAGE in relation to profiles of suicidal ideation, psychological distress and wellbeing in a community cohort of adolescents. Three distinct mental health profiles emerged from the latent profile analysis. While the hypothesis that elevated distress and suicidality would be associated with older-appearing brains was partially supported, the pattern of these findings was more nuanced than anticipated. Only the *moderate distress with lower SI* cluster showed significantly higher BrainAGE relative to the *low distress* cluster, with no significant difference observed between the *moderate distress with greater SI* and *low distress* clusters. This effect was consistent across the primary linear regression and the secondary longitudinal model using all repeated observations. However, when the analytic sample was extended to include observations up to 17.9 years, this cluster effect was no longer evident. Thus, this pattern between suicidality severity and BrainAGE is a central finding that warrants careful consideration.

### 4.1 LPA profile characteristics

The *low distress* cluster was characterised by high wellbeing, low psychological distress and suicidality, and the lowest within-person variability across all four indicators. The *moderate distress with greater SI* cluster exhibited the most severe profile overall, with the lowest wellbeing, and the highest levels of psychological distress and suicidality. This cluster also demonstrated the greatest within-person variability in suicidality scores, suggesting episodic rather than stable suicidality. This pattern is consistent with the fluctuating nature of suicidality (49). The *moderate distress with lower SI* cluster occupied a somewhat intermediate position on most indicators, with elevated psychological distress, lower suicidality and intermediate wellbeing. Within-person variability was also moderate across indicators, suggesting a more stable, elevated distress profile without prominent suicidal features.

### 4.2 BrainAGE observations

Contrary to the hypothesis, the *moderate distress with greater SI* cluster was not found to show advanced brain-ageing. However, the *moderate distress with lower SI* cluster demonstrated greater BrainAGE relative to the *low distress cluster*, and the average BrainAGE for all groups was negative over time. This pattern is partially consistent with the existing literature. Having a history of a mental and behavioural disorders has been linked to advanced biological ageing (10), however such generalisation is not as conclusive as decelerated brain-ageing has been reported in cohorts of generalised anxiety disorder and mood disorder cohorts (24, 25). Taken together, these findings suggest that distinct psychopathological profiles may be associated with different patterns of neurodevelopmental maturation, rather than uniformly advancing ageing as a function of distress severity. Importantly, the association between the *moderate distress with lower SI* cluster and advanced BrainAGE remained significant after controlling for chronological age and biological sex, suggesting that these differences were not solely attributable to demographic differences between clusters.

One possible interpretation is that the *moderate distress with lower SI* cluster reflects a pattern of distress associated with relatively advanced neurodevelopmental maturation, although this explanation remains speculative. Resilience literature suggests that exposure to manageable levels of adversity may promote adaptive neurodevelopment through stress inoculation processes (50, 51). While evidence linking stress inoculation to advanced BrainAGE in adolescents is limited, this framework provides one potential explanation for the observed pattern. Additionally, the distinction between chronic and acute stress profiles may account for the difference in brain-ageing trajectories. The cluster with greater SI had elevated within-person variability, suggesting possible episodic, crisis-level distress. In contrast, the lower suicidal ideation cluster demonstrated less variability in suicidality scores, which may reflect a more stable, chronic pattern of distress over time. Chronic low-grade stress may have cumulative effects on neurodevelopmental trajectories, whereas acute stress may trigger HPA axis dysregulation (12), that disrupts rather than advances maturation. Consistent with de Nooij et al. (24), who found decelerated predicted brain-age among young people with a genetic predisposition to mood disorder who went on to develop one. It also aligns with evidence that severe or clinical-level psychopathology does not always produce the same biological ageing trajectory as subclinical chronic distress (10). Acute or clinical-level distress may disrupt rather than accelerate maturation. Given the instability of this cluster effect when the sample was extended beyond 16.9 years, these interpretations should be considered cautiously and replicated in other longitudinal adolescent samples.

Lastly, the CentileBrain pipeline is a normative dataset and the training data that built the model may not adequately represent adolescent populations with elevated distress or clinical symptomology. Furthermore, this pipeline does not calculate relative contributions of brain regions, nor consider that regional brain development in adolescence follows an uneven, non-linear trajectory (7, 9). This consideration is important for research exploring the relationship between distress profiles among developing cohorts, as advanced maturation of specific regions may be offset by delayed maturation of other regions (5, 9). This could have influenced the findings that the *moderate distress with lower SI* cluster appeared to have more advanced brain aging than the *moderate distress with greater SI* cluster, though this is not conclusive and the small sample sizes should not be ignored.

Post hoc analyses suggested that the cluster-BrainAGE association was not driven by sex adjustment, as removing sex as a covariate did not materially change cluster effects. Removing participant as a random intercept yielded similar findings for the *moderate distress with lower SI* cluster, indicating the finding was not sensitive to the inclusion of a random intercept. When the secondary longitudinal model was extended to the full 12-17.9-year sample, the previously elevated BrainAGE in the *moderate distress with lower SI* cluster was no longer significant. In contrast, chronological age remained a significant positive predictor of BrainAGE across clusters. Overall, these findings suggest that the observed association between mental health profile and BrainAGE may be unstable across small extensions of the age range and sensitive to sample composition or modelling choices. This may reflect developmental changes in sex- and age-related variance or shifts in cluster membership with the inclusion of older and more varied, observations. Results should be cautiously interpreted, and this instability is an important finding for both interpretation and future work.

Although biological sex was not a significant predictor of BrainAGE in the present study, the analysis may have been underpowered to detect sex-specific effects, particularly given the small size of the two moderate distress clusters. Evolutionary-developmental accounts propose sex-specific selection pressures on the timing of maturation, with females advancing and males potentially delaying development under conditions of stress and environmental instability (9). Thus, future research should examine whether the relationship between mental health profiles and BrainAGE differs by sex, including potential sex-by-cluster interactions, as previous studies have reported associations between sex, age, and biological age (9, 18).

### 4.3 Limitations and future directions

While a strength of this study was its use of validated self-report measures (35, 36, 40, 41) and its use of a publicly available, established brain-age estimation tool (6, 42), several limitations should be noted. First, the LPA used participants’ mean and standard deviation scores across time, allowing overall levels and variability to be captured; however, this approach reduced the temporal richness of the longitudinal dataset. Second, while a person-centred cluster analysis was used and the minimal sample size was achieved for each cluster (48), the sample size of both moderate distress clusters was far smaller than the low distress cluster, limiting the ability to examine moderators such as sex or age. Third, this study did not examine bidirectional effects of PBA and mental health profiles over time. Future studies should employ statistical methods to examine directional effects. Fourth, this study did not assess pubertal stage, which is relevant given the evidence that stress effects on biological maturation may be more pronounced at early Tanner stages (18). Future studies should incorporate pubertal measures to better characterise the developmental context of brain-ageing trajectories. Fifth, the mental health profile-BrainAGE association was not stable when extended to 17.9 years, underscoring the need for replication.

The current study identified three distinct mental health profiles in a community cohort of adolescents. *The moderate distress with lower SI* group exhibited significantly higher BrainAGE than the *low distress* reference group, even after controlling for chronological age and biological sex. While this pattern was replicated in the analysis in the secondary longitudinal analysis it was not supported when the sample was extended to include participants age 17-17.9. Contrary to the original hypothesis, the profile characterised by the highest suicidality burden did not differ significantly from the *low distress* group in BrainAGE. The present study lays an empirical foundation for the field to expand on. While there are some suggested interpretations of the mechanisms driving these findings, further longitudinal research is required to understand how wellbeing-distress-suicidality profiles are associated with BrainAGE across adolescence, consider the relative contributions of specific brain regions, and clarify the directional relationships between distress profiles.

## Acknowledgments

Thank you to the young people who generously time to participate in this research. Thank you also to the Thompson Institute radiographers and LABS research assistants who assisted with data collection, Pete Embleton who was instrumental in successfully processing the MRI data on the UniSC HPC, and to Monica D Rosenberg for reviewing the manuscript and providing suggestions based on your expertise. The authors acknowledge the use of Claude Sonnet 5 and Microsoft 365 Copilot (bizchat.20260701.51.4) to assist with code development, syntax refinement, and troubleshooting during the data analysis process. All analytical decisions, interpretation of results, and manuscript content were reviewed and verified by the authors, who take full responsibility for the work.

## Author contribution statements

Maddison Crethar: methodology, formal analysis, investigation, data curation, writing – original draft, visualization; Amanda Boyes: conceptualisation, methodology, investigation, data curation, writing – review and editing, supervision, project administration, funding acquisition; Laura K M Han: conceptualisation, methodology, writing – review and editing, visualization, supervision; Taliah Prince: investigation, writing – review and editing. Lia Mills: investigation, writing – review and editing; Timothy J Silk: writing – review and editing; Nandita Vijayakumar: writing – review and editing; Daniel Hermens: conceptualisation, methodology, writing – review and editing, project administration, resources, funding acquisition, supervision.

## Data availability

The datasets generated and analysed during the current study are not publicly available due the fact that they constitute an excerpt of research in progress but are available from the corresponding author on reasonable request.

