## Supplemental Table 2 for "Adolescent brain-age differences between profiles based on suicidal ideation, distress and wellbeing"

*Supplementary Material S2*

| Predictor | *B* | SE | p |
| --- | --- | --- | --- |
| Intercept | **-13.02** | **0.88** | ***p* <.001** |
| *Moderate distress with greater SI* vs *Low distress* | -0.85 | 0.67 | *p* = .206 |
| *Moderate distress with lower SI* vs *Low distress* | -0.75 | 0.59 | *p* = .210 |
| Sex (female vs male) | 0.76 | 0.44 | *p* =.091 |
| Chronological age | **0.72** | **0.06** | ***p* <.001** |

Post hoc comparisons against the secondary, longitudinal analysis (Linear mixed model; BrainAGE ~ cluster + sex + age + (1 | ParticipantID)) using the 12-17.9 dataset. Cluster categorisation remained similar however the distribution across clusters differed. *Low distress, N=*97 with 538 observations; *Moderate distress with lower SI, N=*23 with 140 observations; *Moderate distress with greater SI, N=*17 with 72 observations.
