## Supplemental Table 1 for "Adolescent brain-age differences between profiles based on suicidal ideation, distress and wellbeing"

**Supplementary material**

*Supplementary Material S1*

| Predictor | Linear mixed model | Fixed-effects model  (random intercept excluded) | Linear mixed model  (sex excluded) |
| --- | --- | --- | --- |
| Intercept | **-12.39 (0.98), *p*<.001** | **-7.14 (1.18), *p*<.001** | **-12.25 (0.97), *p*< .001** |
| *Moderate distress with greater SI* vs *Low distress* | -1.19 (0.68), *p=*.083 | -1.00 (0.41), *p*=.016 | -1.14 (0.68), *p*=.095 |
| *Moderate distress with lower SI* vs *Low distress* | **1.87 (0.76), *p*=.015** | **1.76 (0.39), *p*<.001** | 2.02 (0.74), *p*=.007 |
| Sex (female vs male) | 0.37 (0.42), *p*=.380 | 0.68 (0.22), *p*=.002 | - |
| Chronological age | **0.67 (0.07), *p*=<.001** | **0.28 (0.08), *p*<.001** | **0.68 (0.067), *p*< .001** |

Post hoc comparisons against the secondary, longitudinal analysis (Linear mixed model; BrainAGE ~ cluster + sex + age + (1 | ParticipantID)), linear mixed model with no random intercept (model: BrainAGE ~ cluster + sex + age) and linear mixed model without sex (model: BrainAGE ~ cluster + age + (1 | ParticipantID)). The dataset for each exploratory post hoc remained restricted to 12-16.9 years.
